# Motor Learning And Savings Of Adaptive Mediolateral Control During Split-Belt Walking

**DOI:** 10.1101/2024.12.29.630661

**Authors:** Norah M. Nyangau, Alysha T. Bogard, Aviva K. Pollet, Logan M. Pellegrino, Andrew Q. Tan

## Abstract

Active control of frontal plane mechanics regulates balance in destabilizing environments, such as during asymmetric split-belt walking. Compared to sagittal plane mechanics, mediolateral (ML) kinematic and kinetic adaptations to split-belt perturbations are not as extensively reported. Moreover, the associated metabolic cost of these adaptations as well as the retention of previously learned ML adaptations upon re-exposure to the same perturbation have not been concurrently examined. We investigated adaptations in step width and peak ML ground reaction forces (GRF) during an initial and subsequent perturbation in order to characterize motor learning and motor savings, respectively. Additionally, we examined the extent to which a neuroplasticity inducing stimulus, acute intermittent hypoxia (AIH), affected the magnitude of each adaptation. Although we observed bilateral increases in step width during the initial adaptation, only the slow leg significantly reduced step width during the subsequent perturbation. Distinct interlimb differences emerged as only the slow leg modulated ML GRF during the braking phase whereas the fast leg increased ML GRF during the propulsive phase. The AIH group uniquely demonstrated greater motor savings of reduced step width and peak ML GRF strategies during the propulsive phase, suggesting greater retention of prior strategies. Furthermore, we find significant associations between ML kinetic adaptations and reductions in metabolic cost. Together, our findings suggest that unlike the sagittal plane, asymmetrical frontal plane adaptations contribute to ML stability as well as reductions in metabolic cost during split-belt walking. These insights could inform clinical training approaches to improve balance and prevent falls in clinical populations.

**NEW & NOTEWORTHY:** We investigated adaptations in step width and mediolateral ground reaction forces during the braking and propulsive phases of split-belt walking across an initial and subsequent perturbation. We observe that the initial learning and savings of unique interlimb frontal plane coordination strategies contribute to stability and are associated with a reduction in metabolic cost.

## INTRODUCTION

Maintaining balance during walking is critical for preventing falls, especially in older adults (Rogers et al., 2001; McIlroy & Maki, 1996) and individuals with neurological impairments, such as spinal cord injury (Jørgensen et al., 2016; Arora et al., 2019). In contrast to predominantly passive control of sagittal plane mechanics (McGeer, 1990; Kuo & Donelan, 2010), it is well documented that active control of frontal plane mechanics is required to maintain mediolateral (ML) stability (Bauby & Kuo, 2000; Donelan et al., 2004; Kuo & Donelan, 2010). To enhance lateral margin of stability, measured as the minimum distance between the base of support and extrapolated center of mass (Hof et al., 2005), able-bodied individuals often regulate foot placement (Bruijn & Van Dieën, 2018; Buurke et al., 2018; Rawal & Singer, 2021) to widen their stance (McAndrew Young & Dingwell, 2012; Hak et al., 2013). Indeed, individuals with neurological impairments often adopt wider strides compared to healthy controls (Curtze et al., 2024). While spatiotemporal and kinetic features of sagittal plane mechanics have been extensively studied to characterize balance during gait (Park & Finley, 2017; Debelle et al., 2020), adaptations in frontal plane kinetics are less comprehensively reported. Particularly, the relationship between adaptive step kinematics and their associated ML ground reaction forces (GRF) during sustained destabilizing perturbations has not been thoroughly examined. Given that both ML foot placement and GRF are coupled responses to external destabilization (Rawal & Singer, 2021), characterizing their simultaneous adaptations is necessary to further elucidate control priorities in frontal plane mechanics.

One approach to assess ML balance control is through split-belt walking, which uses single-belt speed perturbations to induce transient spatiotemporal walking asymmetries (Sombric & Torres-Oviedo, 2020; Sánchez et al., 2021). These perturbations elicit sensorimotor error-based motor learning (Reisman et al., 2005; Roemmich & Bastian, 2015; Leech et al., 2022), as evidenced by progressive reductions in step length asymmetry and double support time asymmetry within the sagittal plane (Donelan et al., 2002; Sánchez et al., 2021). Importantly, participants also demonstrate retention of the learned sagittal plane motor strategies during subsequent exposure to the same perturbation known as motor savings (Roemmich & Bastian, 2015; Leech et al., 2018; Bogard et al., 2023). While frontal plane adaptations have been documented during a single exposure to a split-belt speed perturbation (Buurke et al., 2018, 2019, 2021; Cornwell et al., 2024), motor savings have not been directly examined. Motor savings is thought to reflect the retention of previously learned motor patterns (Leech et al., 2018; Huang et al., 2012), as well as shifts in reactive vs. anticipatory control strategies (Rawal & Singer, 2021; Ahuja & Franz, 2022). Thus, examining the temporal evolution of ML coordination strategies during both the initial and subsequent perturbation exposure could reveal further insights into the adaptive control of balance when stability is challenged.

During split-belt adaptation, indices of metabolic cost, like net metabolic power, accompany learned biomechanical adaptations during both motor learning and motor savings (Finley et al., 2013; Sánchez et al., 2017; Price et al., 2023). Notably, we and others have demonstrated that net metabolic power initially increases (Sánchez et al., 2019; Butterfield & Collins, 2022; Bogard et al., 2023) but gradually decreases toward baseline values as participants adopt more energetically efficient coordination strategies. In the sagittal plane, reductions in net metabolic power in able-bodied individuals parallel improvements in spatiotemporal symmetry during split-belt adaptation (Finley et al., 2013; Bogard et al., 2023). These findings suggest that energy optimization may play a role in split-belt adaptation (Emken et al., 2007; Sánchez et al., 2017; Stenum & Choi, 2020) as different control strategies exact distinct metabolic demands (Ahuja & Franz, 2022).

In contrast with the sagittal plane, concurrent reductions in net metabolic power and adaptations in frontal plane mechanics have been inconsistently observed across split-belt walking trials. For example, while external ML stabilization has been estimated to reduce metabolic cost during normal walking (Donelan et al., 2004; Dean et al., 2007), changes in ML margin of stability or step width during split-belt walking show no consistent relationships with metabolic cost (Buurke et al., 2018). Deviations in step width (Donelan et al., 2004), whole-body angular momentum (Cornwell et al., 2024) and ML foot roll-off (Buurke et al., 2018) suggest that other ML gait parameters influence metabolic cost during split-belt adaptation. To our knowledge, no study has concomitantly characterized frontal plane mechanics and metabolic adaptations during both motor learning and motor savings. Thus, our primary aim was to examine kinematic and kinetic adaptations alongside associated changes in net metabolic power during split-belt walking.

We recently demonstrated that brief exposure to low oxygen, known as acute intermittent hypoxia (AIH), enhances sagittal plane adaptation during split-belt walking, including spatiotemporal and anterior-posterior force asymmetry (Bogard et al., 2023). Although both the AIH and control groups successfully adapted their walking mechanics, the AIH group achieved a greater reduction in net metabolic power (Bogard et al., 2023). Combined with prior studies that show AIH facilitates descending excitability and reduces performance fatiguability (Bogard et al., 2024; Bogard, Pollet, et al., 2024), these observations suggest that adaptive changes within the nervous system may facilitate greater motor learning. Indeed, AIH-induced synthesis of brain-derived neurotrophic factor (BDNF) parallel strengthened synaptic plasticity and improvements in motor performance in rodents (Fritsch et al., 2010; Lovett-Barr et al., 2012). Increases in BDNF in humans have also been associated with enhanced motor learning (Rasmussen et al., 2009; Leech & Hornby, 2017). Thus, to further elucidate the neural control process underlying motor adaptation, a secondary aim of this study was to investigate whether repetitive AIH enhances motor learning of frontal plane adaptation and whether such changes drive greater reductions in net metabolic power.

Accordingly, we examined time-dependent adaptations in step width and peak ML GRF during braking and propulsive phases of gait in response to split-belt speed perturbations. We further characterized the extent to which these adaptations are retained in response to a second exposure to the same perturbation, as well as the corresponding effect on net metabolic power. We hypothesized that participants would initially adopt a wider step and greater ML GRF and that these coordination strategies would be reduced upon subsequent exposure. Given that AIH has been shown to improve sensorimotor adaptation in sagittal plane mechanics (Bogard et al., 2023), we tested the secondary hypothesis that adaptations in frontal plane mechanics and reductions in net metabolic power would be more prominent in the AIH group relative to the control group.

## MATERIALS AND METHODS

### Participants

Thirty able-bodied individuals with no prior history of neurological impairments were recruited for the study. All participants provided informed consent approved by the Colorado Multiple Institutional Review Board (COMIRB no. 20-0689). The procedures complied with the standards of the *Declaration of Helsinki* and the study was registered on clinicaltrials.org (NCT05341466). Participants were randomized into an AIH group (*n* = 15, 9 females, 6 males, age 23.5 ± 2.3 years, body mass 65.9 ± 11.5 kg) or a control group (*n* = 15, 7 females, 8 males, age 25.3 ± 5.3 years, body mass 73.9 ± 15.1 kg). Criteria for exclusion included prior exposure to split-belt walking, altitude sensitivity, cardiovascular or pulmonary diseases, syncope, or being pregnant at the time of the study.

### Protocols

#### Acute Intermittent Hypoxia

For the AIH group, low-oxygen air was delivered through an altitude generator (HYP 123; Hypoxico, Inc., USA) while trained researchers continuously monitored heart rate (HR) and oxygen saturation levels (SpO2) along with measuring blood pressure (BP) every 5 normoxic cycles (Masimo; Irvine, CA, USA). Similar to Tan et al., 2021, AIH consisted of 90 s bouts of hypoxic air (9% O2) followed by 60 s bouts of normoxic air (21% O2) for a total of 15 cycles. Treatment was paused if individuals de-saturated below 70% SpO2 and resumed when they re-saturated above 80% (Tan et al., 2020). The experiment was discontinued if any of the following conditions occurred: systolic BP exceeded 140 mmHg, diastolic BP exceeded 90 mmHg, or HR values surpassed 160 bpm; however, these thresholds were not exceeded (Tan et al., 2020). Termination criteria also included reported or visible signs of adverse events such as dizziness, numbness, tinnitus, blurred vision, or diaphoresis. This protocol was repeated for five consecutive days at approximately the same time each day. All participants tolerated the hypoxic dose and completed the full procedure.

#### Gait Mechanics

Kinematic data was recorded with a 10-camera system at a rate of 100 Hz (Vicon Nexus v2.8.1; Vicon Motion Systems, Denver, CO, USA). Thirty-four reflective markers were placed as shank and thigh clusters and on the following anatomical landmarks: anterior and posterior superior iliac spines, iliac crests, greater trochanters, medial, and lateral femoral epicondyles, medial and lateral malleoli, calcanei and first and fifth metatarsals (Montgomery & Grabowski, 2018). Participants were instructed to avoid using the handrails unless necessary for safety (Buurke, Lamoth, Van Der Woude, & Den Otter, 2019) and were secured with a single passive harness that neither affected movement nor provided body weight support. A mirror was positioned in front of the treadmill to help individuals avoid crossing their feet onto the opposite belt. Time-synchronized kinetic variables were measured on an instrumented split-belt treadmill at a rate of 1000 Hz, and the belt speeds were controlled independently (M-Gait, D-flow v3.34.3; Motek Medical, Houten, NL).

#### Metabolic Data

Metabolic rate was measured using an open circuit spirometry system (TrueOne 2400; ParvoMedics Inc., Salt Lake City, UT, USA). The rate of oxygen consumption (*V̇*_0_2__) and carbon dioxide production (*V̇*_C0_2__) was utilized to calculate respiratory exchange ratios (RER = *V̇*_C0_2__/*V̇*_0_2__). All participants had RER values below 1, indicating that aerobic pathways were primarily being used (Huang et al., 2012). Resting metabolic rate (RMR) was estimated from each individual’s average *V̇*_0_2__ and *V̇*_C0_2__ during the last 2 minutes of a 5-minute standing trial for the AIH (1.51 ± 0.26 W/kg) and control (1.50 ± 0.20 W/kg) groups. We estimated energetic cost during walking trials by calculating metabolic power using the regression formula provided in Equation 1 (Péronnet & Massicotte, 1991).

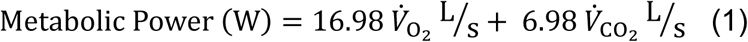

Net metabolic power (W/kg) was obtained from the difference between metabolic power and average RMR and normalized to body mass (Finley et al., 2013). Data from the initial 60 s of each trial were excluded, as this reflects the duration needed for net metabolic power to stabilize to its average value (Finley et al., 2013).

#### Split-Belt Walking Protocol

We followed the split-belt walking protocol described by Bogard et al., 2023 which included four trials of two tied-belt and two split-belt walking conditions. A single belt was randomly selected to speed up without forewarning (Reisman et al., 2005) throughout the experiment. The first trial was ‘baseline,’ which included walking at a tied-belt speed of 1.0 m/s for 300 strides. Following baseline, participants performed an ‘adapt 1’ trial that involved tied-belt walking at 1.0 m/s for 15-30 strides before a sudden increase of one belt to 2.0 m/s for 300 strides of split-belt walking at a 2:1 belt speed ratio. Next, the participants performed a ‘washout’ trial where they walked with tied-belt speed of 1.0 m/s for 350 strides. Finally, participants performed an ‘adapt 2’ trial by walking at 1.0 m/s tied-belt speed for 15-30 strides followed by a second exposure to the split-belt perturbation at a 2:1 belt speed ratio for 300 strides. The trials were performed consecutively, stopping after each trial with an optional 1-minute break during which individuals were instructed to stand still (Leech et al., 2018; Sombric & Torres-Oviedo, 2020). Participants in the intervention group performed this protocol 15 minutes after receiving their final low-oxygen treatment.

### Data Analysis

Motion-captured markers were labeled in Vicon to create lower body models for gait analysis using custom pipelines created in Visual3D (v2021.11.3; HAS-motion Inc., Germantown, MD, USA). Ground-reaction forces (GRF) were filtered using Lowpass Butterworth with a cutoff frequency of 20 Hz. Heel-strike and push-off gait events were identified at vertical GRF thresholds of 30 N (Karakasis & Artemiadis, 2021). Step widths were calculated in Visual3D as the perpendicular distance between a stride vector formed by consecutive heel-strikes of the contralateral limb and the calcaneus position at heel-strike of the ipsilateral limb. Midstance events were defined as the instance when anterior-posterior GRF crossed zero when plotted, marking the transition from braking into propulsion phase of gait (Masani et al., 2002). Therefore, the time intervals between heel-strike and midstance defined the braking phases, and the propulsive phases were defined between midstance and push-off. We identified peak ML GRF during both the braking and propulsive phases within each step.

Step width and peak ML GRF values were averaged for each trial. The last 20 steps of the baseline were analyzed for comparison with the perturbation trials. The perturbation trials were divided into separate learning phases. Namely, ‘early learning’ was defined as the first 5 steps immediately following the change in belt speed while ‘late learning’ was the last 20 steps of each trial. The early and late learning phases within a single perturbation trial are referred to as motor learning. To characterize the retention of motor strategies, we compared the learning phases across perturbation trials (Bogard et al., 2023). ‘Early savings’ compared motor adaptions between early adapt 1 and early adapt 2, while ‘late savings’ compared motor adaptations between late adapt 1 and late adapt 2. Comparisons of learning phases across perturbation trials are described as motor savings.

### Statistical Analysis

Kinematic, kinetic, and metabolic data were processed in MATLAB (R2023b; The MathWorks, Inc., Natick, MA, USA). Statistical analysis was performed in R Studio (v2024.04.2) with the significance set to *p* < 0.05. Normality and homogeneity of the data were tested using Shapiro-Wilks and Levene’s tests, respectively. An *a priori* power analysis using preliminary data determined that a sample size of 14 participants per group would be adequate to observe differences between and within groups (power = 0.85, Cohen’s *f* = 0.76, *α* = 0.05, *F*(1,12) = 4.74) (Cohen, 1988). Two-way repeated measures analysis of variance (ANOVA) tests were utilized to examine the effect of leg speed on step width. Mean step width was compared between the fast and slow legs at baseline, early adapt 1, late adapt 1, early adapt 2, early adapt 2, and late adapt 2. Similarly, two- way ANOVAs were conducted to examine the effect of leg speed on peak ML GRF. Peak ML GRFs were compared between the fast and slow legs during the braking and propulsive phases at baseline, early adapt 1, late adapt 1, early adapt 2, and late adapt 2. Furthermore, two-way ANOVAs were conducted separately for each limb to investigate the effects of AIH intervention on early and late savings of step width and peak ML GRF. Additional two-way ANOVAs analyzed the effects of motor learning and motor savings on net metabolic power. Average net metabolic power was compared between early and late learning of adapt 1 and 2 as well as early and late savings. For analyses of repeated measures that violated the assumption of sphericity (*P* < 0.05) as determined by Mauchly’s Test for Sphericity, adjustments were made using the Greenhouse-Geisser (GG) correction for epsilon (GGe) values below 0.75 or the Huynh-Feldt (HF) correction for GGe values above 0.75. Tukey’s Honestly Significant Difference (HSD) tests were performed to identify pairwise interactions for main effects and interactions that reached significance. Linear regression analyses were generated to investigate relationships between changes in net metabolic rate and changes in step width and peak ML GRF.

## RESULTS

### Kinematic Adaptations

#### Reductions in Step Width During Motor Learning

During adapt 1, learning phase had a main effect on step width (*F*(1.38, 39.9) = 18.2, *P* < 0.001), in which both legs increased step width during early learning then decreased in late learning toward baseline levels (Fig. 1*A*). Post-hoc comparisons revealed significant increase from baseline to early learning (fast leg: *t*(60.4) = -5.35, *P* < 0.001; slow leg: *t*(60.4) = -5.14, *P* < 0.001) and decrease from early learning to late learning (fast: *t*(60.4) = 5.18, *P* < 0.001; slow: *t*(60.4) = 5.03, *P* < 0.001). No main effects of leg speed (*P* = 0.491) nor an interaction of leg speed and learning phase (*P* = 0.749) on step width were observed.

**Figure 1.**
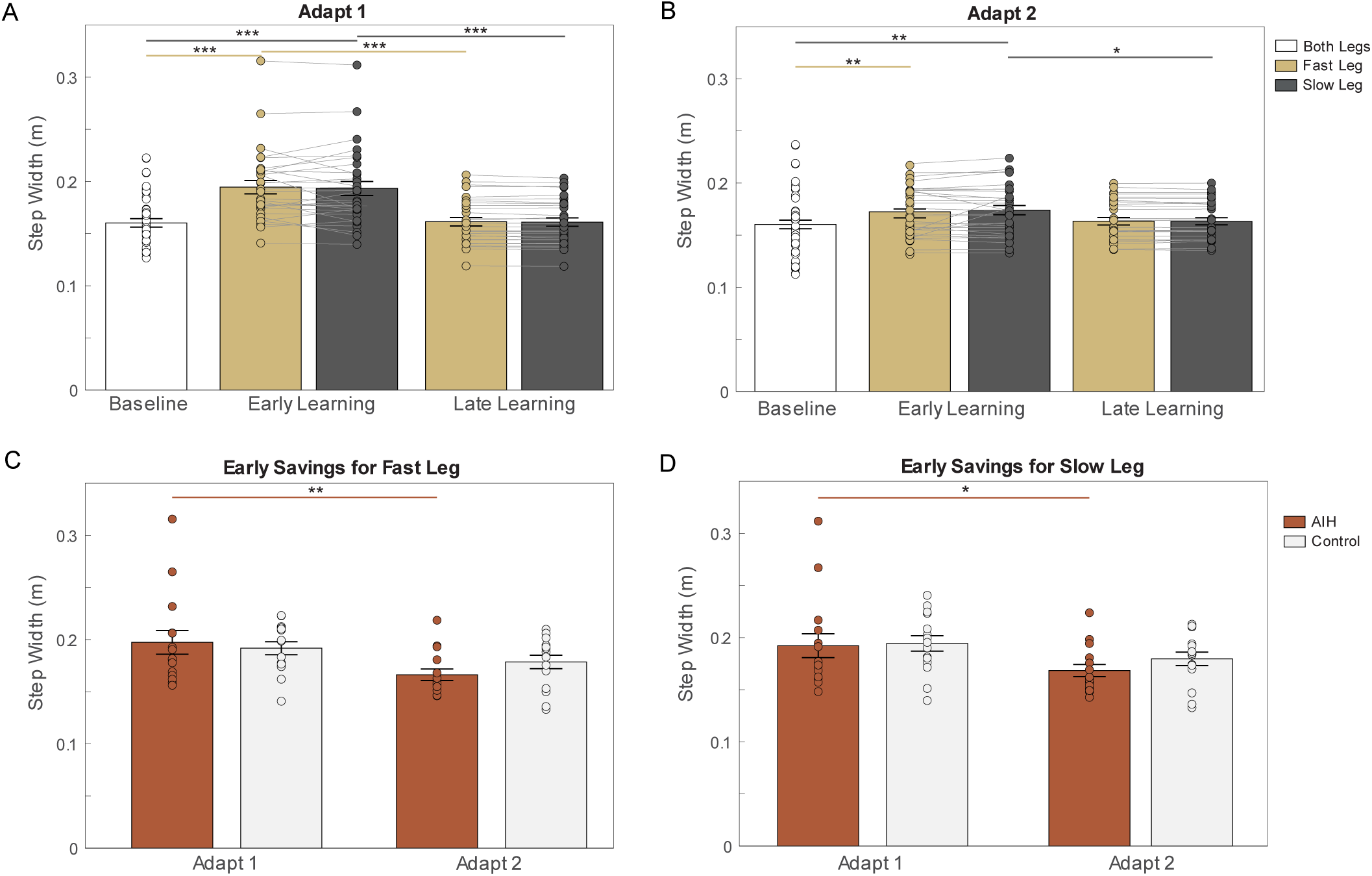
Step width adaptation. *A*) Both legs increased their step width from baseline during early learning then significantly narrowed during late learning of adapt 1. *B*) During adapt 2, both legs widened their steps during early learning but only the slow leg significantly decreased its width during late learning. The AIH group demonstrated early savings of step width in both the fast leg (*C*) and slow leg (*D*) compared to controls. Bar graphs represent mean step width and standard error (SE). ∗ *P* < 0.05, ∗∗ *P* < 0.01, ∗∗∗ *P* < 0.001.

Similarly, step width significantly narrowed throughout learning during adapt 2 (*F*(2, 58) = 6.19, *P* = 0.004), but neither leg speed (*P* = 0.348) nor an interaction of leg speed and learning phase (*P* = 0.482) had significant effects (Fig. 1*B*). Pairwise *t*-tests showed significantly larger widths during early learning compared to baseline for both legs (fast leg: *t*(62.1) = -3.12, *P* = 0.008; slow leg: *t*(62.1) = -3.49, *P* = 0.002). Interestingly, differences between limbs began to emerge during late learning of the second perturbation, with significant narrowing observed in the slow leg (*t*(62.1) = 2.74, *P* = 0.021).

#### Reductions in Step Width During Motor Savings

To examine how step width adapts independently for each limb, we analyzed the fast and slow limbs separately, highlighting motor savings strategies of AIH and control groups. For the fast leg, we observed reduced step widths as a main effect of perturbation between early learning phases (*F*(1, 28) = 13.4, *P* = 0.001), but no effect of group (*P* = 0.717) nor an interaction of perturbation and group (*P* = 0.149) (Fig. 1*C*). Pairwise comparisons identified early savings strategies in the AIH group (*t*(28) = 3.64, *P* = 0.001) but not the controls (*P* = 0.013). We observed no late savings on the fast leg between perturbation (*P* = 0.481), within groups (*P* = 0.053), nor the interaction of these factors (*P* = 0.123), indicating that participants in both groups had adapted by this timeframe.

Similarly, a repeated measures ANOVA for the slow leg revealed reductions in step width as a main effect of perturbation during early learning phases (*F*(1, 28) = 9.27, *P* = 0.005) (Fig. 1*D*). Post-hoc revealed early savings of smaller step widths in the AIH group (*t*(28) = 2.66, *P* = 0.013) but not the control group (*P* = 0.111). There were no late savings across perturbations (*P* = 0.403), within groups (*P* = 0.606), nor an interaction of either (*P* = 0.120).

### Kinetic Adaptations

#### Braking Phase During Motor Learning

Peak ML GRF was significantly larger than baseline during early learning for both legs then decreased for just the slow leg during late learning (Fig. 2*A*). We observed main effects of leg speed (*F*(1,29) = 47.2, *P* < 0.001), learning phase (*F*(1.40,40.8) = 61.4, *P* < 0.001), and interaction between leg speed and learning phase (*F*(1.66,48.2) = 34.0, *P* < 0.001) for peak ML GRF during the braking phase of adapt 1. Post-hoc analyses showed a significant increase for the fast leg between baseline and early learning (*t*(93.1) = -9.96, *P* < 0.001) and between baseline and late learning (*t*(93.1) = -9.90, *P* < 0.001). The slow leg similarly increased peak ML GRF between baseline and early learning (*t*(93.1) = - 8.95, *P* < 0.001) and decreased from early to late learning (*t*(93.1) = 6.67, *P* < 0.001). Furthermore, we observed significant reductions in force in the slow leg compared to the fast leg during late learning (*t*(86.9) = 10.7, *P* < 0.001).

**Figure 2.**
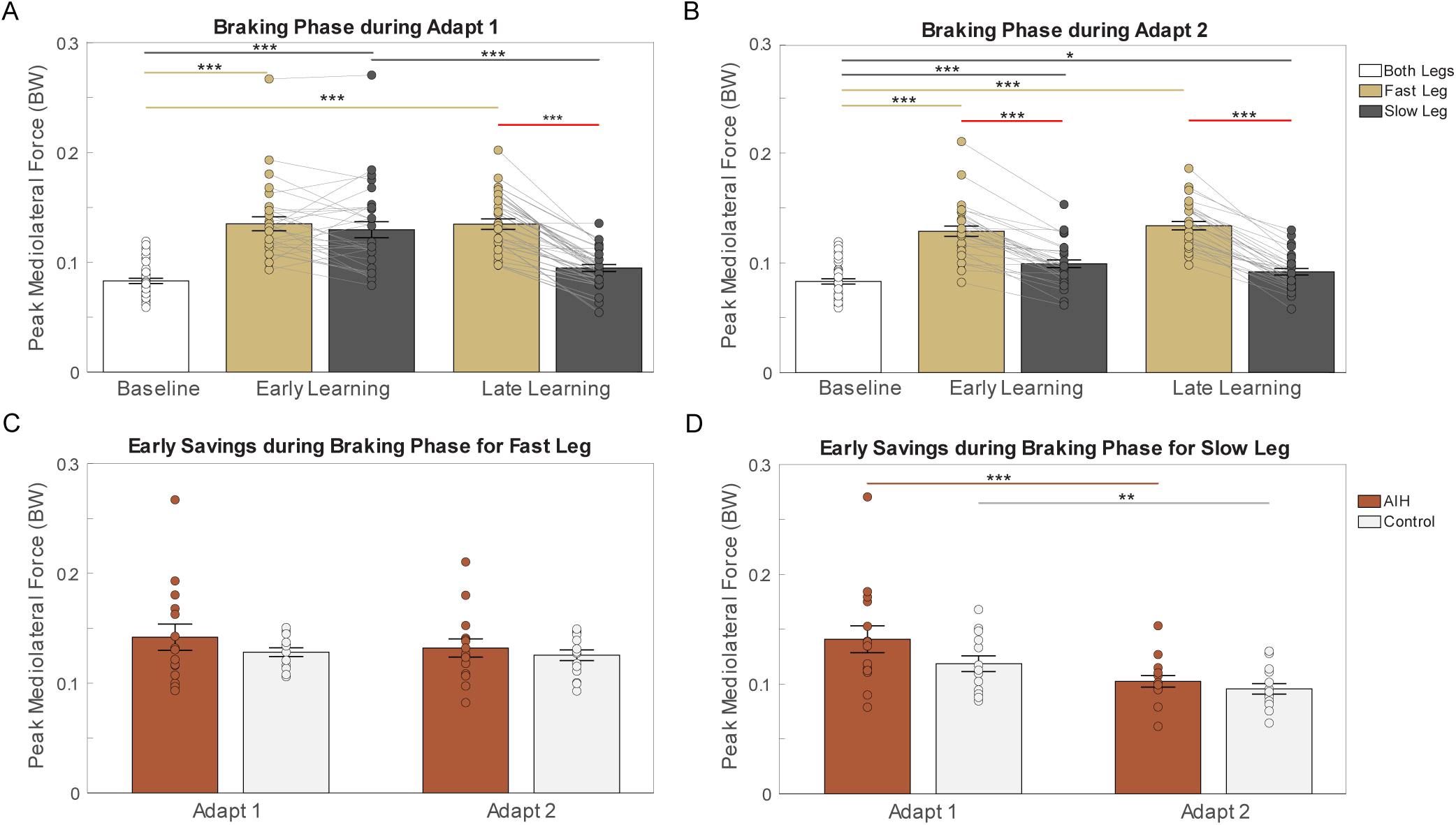
Peak mediolateral ground reaction forces (ML GRF) during braking phase. *A*) peak ML GRF increased from baseline for both legs in early learning and decreased for the slow leg in late learning of adapt 1. *B*) between-limb differences show higher peak ML GRF maintained by the fast leg during early and late learning of adapt 2 compared to the slow leg. There were no early savings strategies observed for the fast leg (*C*), whilst the slow leg significantly decreased magnitude responses during the second adaptation for both groups (*D*). Bar graphs show averaged peak ML GRF magnitudes and standard error (SE). ∗ *P* < 0.05, ∗∗ *P* < 0.01, ∗∗∗ *P* < 0.001. The red line represents significant differences between limbs.

During adapt 2 there were main effects of leg speed (*F*(1,29) = 192, *P* < 0.001), learning phase (F(2,58) = 85.9, *P* < 0.001), and an interaction between leg speed and learning phase (*F*(2,58) = 83.8, *P* < 0.001) (Fig. 2*B*). Post-hoc *t*-tests found significant increase between learning phases for the fast leg (baseline vs. early learning: (*t*(97.1) = -14.5, *P* < 0.001); baseline vs. late learning: (*t*(97.1) = -16.12, *P* < 0.001)) and the slow leg (baseline vs. early learning: (*t*(97.1) = -5.11, *P* < 0.001); baseline vs. late learning: (*t*(97.1) = -2.81, *P =* 0.016). Additionally, the fast leg maintained significantly higher peak force magnitudes compared to the slow leg during early learning (*t*(82.2) = 11.5, *P* < 0.001) and late learning (*t*(82.2) = 16.3, *P* < 0.001).

#### Braking Phase During Motor Savings

To examine motor savings strategies between AIH and control groups, peak ML GRF of the fast and slow legs were independently analyzed. For the fast leg, we observed no main effects of group (early savings: *P* = 0.332; late savings: *P* = 0.965), perturbation (early savings: *P* = 0.165; late savings: *P* = 0.755), nor interaction of group and perturbation (early savings: *P* = 0.428; late savings: *P* = 0.658) (Fig. 2*C*). This suggests that the fast leg’s strategy was to maintain larger peak ML GRF throughout adaptation. Conversely, the slow leg showed main effects of perturbation (*F*(1,28) = 42.1, *P* < 0.001), but no group effect(*P* = 0.164) nor interaction of group and perturbation (*P* = 0.113) for early savings (Fig. 2*D*). Pairwise comparisons showed that both groups demonstrated early savings strategies (AIH: *t*(28) = 5.75, *P* < 0.001); controls: (*t*(28) = 3.43, *P =* 0.002), reflecting a reduction in response magnitudes upon re-exposure to asymmetric walking. Neither group exhibited late savings for the slow leg (*P* = 0.663) between perturbations (*P* = 0.279), nor an interaction of group and perturbation (*P* = 0.305).

#### Propulsive Phase During Motor Learning

We observed main effects of leg speed (*F*(1,29) = 54.5, *P* < 0.001), learning phase *F*(1.38,40.1) = 32.2, *P* < 0.001), and an interaction of leg speed and learning phase (*F*(1.44,41.9) = 65.8, *P* < 0.001) for peak ML GRF during the propulsive phase of adapt 1. The fast leg significantly increased force magnitudes during early learning (*t*(93.5) = - 11.11, *P* < 0.001) which then decreased during late learning (*t*(93.5) = 10.78, *P* < 0.001) (Fig. 3*A*). Pairwise analysis revealed a significant difference in magnitude between the legs during early learning (*t*(87) = 13.5, *P* < 0.001), with the fast leg exhibiting larger magnitudes while the slow leg maintaining forces approaching baseline values.

**Figure 3.**
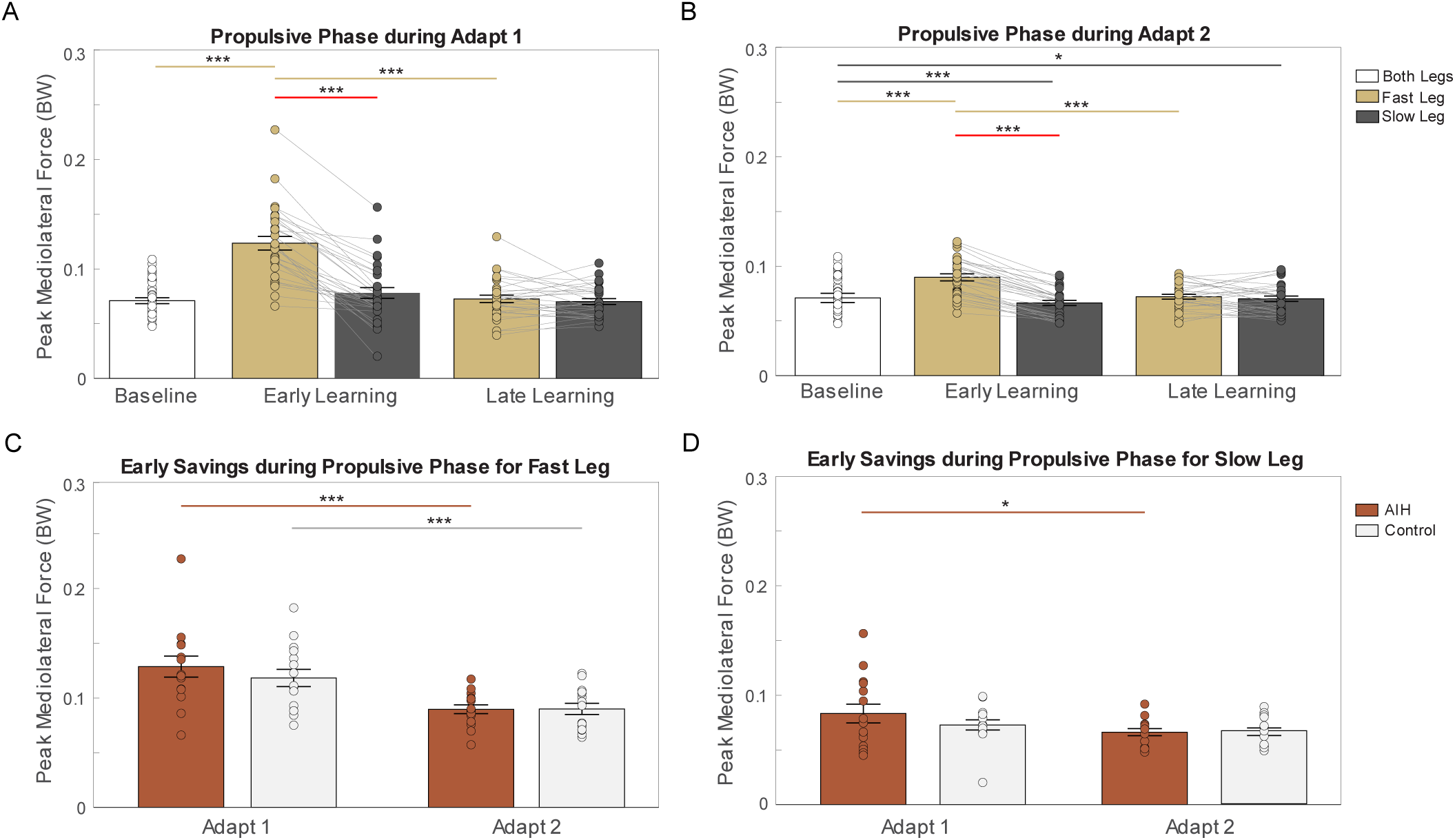
Peak mediolateral ground reaction forces (ML GRF) during propulsive phase. *A*) Only the fast leg increased peak ML GRF during early learning which decreased in late learning of adapt 1 while the slow leg stayed at baseline. *B*) During adapt 2, the fast leg shows a similar adaptation strategy as in adapt 1 and the slow leg decreases peak ML GRF from baseline in early and late learning. *C*) Both the AIH and control groups showed early savings on the fast leg. *D*) Only the AIH group exhibited early savings on the slow leg. Bar graphs show mean peak ML GRF magnitudes and standard error (SE). ∗ *P* < 0.05, ∗∗ *P* < 0.01, ∗∗∗ *P* < 0.001. The red line represents significant differences between limbs.

During adapt 2, we identified main effects of leg speed (*F*(1,29) = 31.7, *P* < 0.001), learning phase (*F*(2,58) = 5.56, *P* = 0.006), and interaction of leg speed and learning phase (*F*(1.62,47.1) = 68.5, *P* < 0.001) (Fig. 3*B*). Pairwise comparisons showed significant increase for the fast leg from baseline to early learning (*t*(95.4) = -7.62, *P* < 0.001) and decrease from early to late learning: (*t*(95.4) = 7.19, *P* < 0.001). The slow leg significantly reduced peak ML GRF below baseline during early learning (*t*(95.4) = 4.01, *P* < 0.001) and late learning (*t*(95.4) = 2.45, *P =* 0.042). Additionally, there were evident interlimb differences during early learning (*t*(85.7) = 12.4, *P* < 0.001), with the fast leg increasing and the slow leg decreasing relative to baseline.

#### Propulsive Phase During Motor Savings

A two-way ANOVA investigated early savings of peak ML GRF on the fast leg during the propulsive phase, which showed a main effect of perturbation (*F*(1,28) = 43.0, *P* < 0.001). There were no effects of group (*P* = 0.559) nor interaction of group and perturbation (*P* = 0.303). Both the AIH (*t*(28) = 5.38, *P* < 0.001) and control (*t*(28) = 3.90, *P* < 0.001) groups demonstrate early savings of reduced peak ML force magnitude (Fig. 3*C*). In late savings, there were no significant main effects of group (*P* = 0.417), perturbation (*P* = 0.833), nor interaction of group and perturbation (*P* = 0.208).

Additionally, an analysis for the slow leg showed a main effect of perturbation (*F*(1,28) = 7.08, *P* = 0.013), but no effect of group (*P* = 0.433) nor interaction of group and perturbation (*P* = 0.225). Post-hoc analysis revealed early savings of peak ML GRF during the propulsive phase, specifically lower peak ML GRF in the AIH group (*t*(28) = 2.76, *P* =

0.010) but not in the controls (*P* = 0.324) (Fig. 3*D*). There were no significant late savings between perturbations (*P* = 0.946), within groups (*P* = 0.852), nor interaction of group and learning phase (*P* = 0.971).

### Metabolic Adaptations

#### Net Metabolic Power During Motor Learning

A two-way ANOVA showed a main effect of learning phase for net metabolic power (*F*(1,28) = 42.48, *P* < 0.001) but no group effect (*P* = 0.903) nor interaction between group and learning phases (*P* = 0.672). Pairwise *t*-tests showed that net metabolic power decreased in late learning of adapt 1 compared to early learning for both the AIH (*t*(28) = 4.91, *P* < 0.001) and control groups (*t*(28) = 4.30, *P* < 0.001) (Fig. 4*A*). During adapt 2, there were main effects of learning phase (*F*(1,28) = 4.50, *P* = 0.043) and interaction of group and learning phase (*F*(1,28) = 5.29, *P* = 0.029) for net metabolic power. Post-hoc comparisons revealed that the AIH group increased net metabolic power during late learning (*t*(28) = -3.13, *P* = 0.004) (Fig. 4*B*).

**Figure 4.**
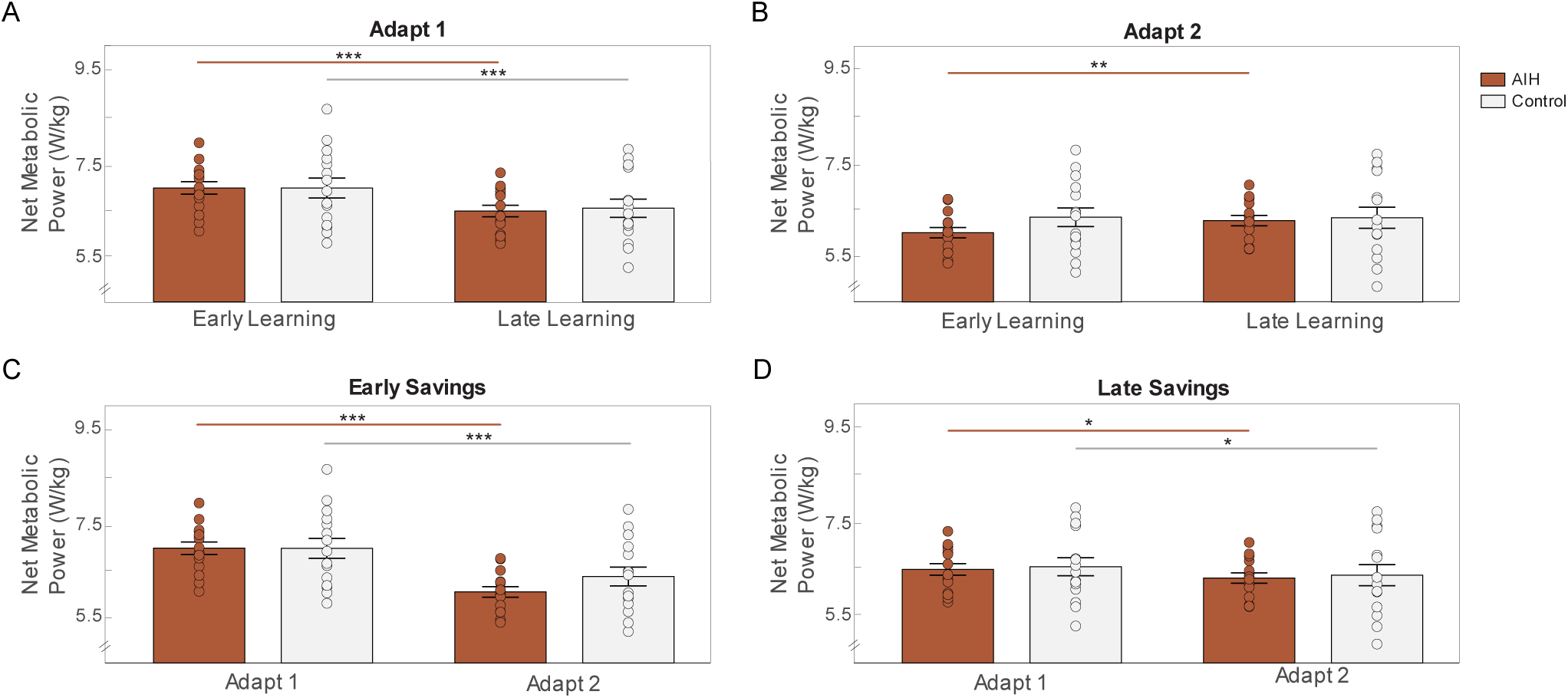
Net metabolic power. *A*) Both groups decreased net metabolic power during late learning of adapt 1. *B*) During adapt 2, the AIH group increased net metabolic power during late learning. Both groups demonstrated early savings (*C*) and late savings (*D*) of net metabolic power. Bar graphs show average net metabolic power (W/Kg) and standard error (SE). ∗ *P* < 0.05, ∗∗ *P* < 0.01, ∗∗∗ *P* < 0.001

**Figure 5.**
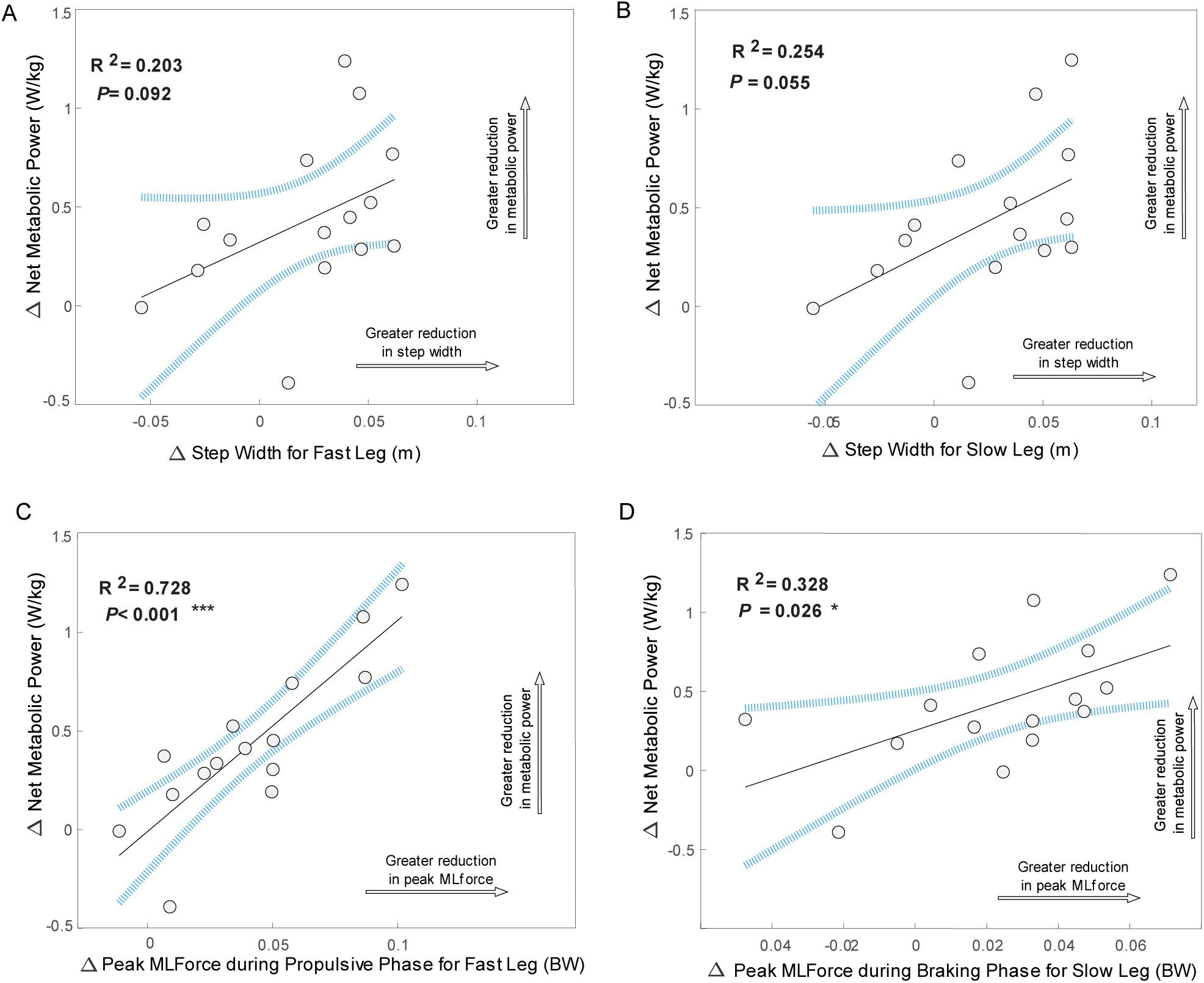
Metabolic regressions for fast and slow legs during adapt 1 in the control group. *A*) Nonsignificant positive relationship between decreases in net metabolic power and step width on the fast leg (R^2^ = 0.203, *P* = 0.092). *B*) Nonsignificant positive relationship between decreases in net metabolic power and step width on the slow leg (R^2^ = 0.254, *P* = 0.055). *C*) Significant positive correlation between decreases in net metabolic power and peak ML GRF during propulsive phase on the fast leg (R^2^ = 0.728, *P* < 0.001). *D*) Significant positive correlation between decreases in net metabolic power and decreases in peak ML GRF during the braking phase on the slow leg (R^2^ = 0.328, *P* = 0.026). Area within blue lines represents 95% confidence and grey points are individual participants’ data. ∗ *P* < 0.05, ∗∗ *P* < 0.01, ∗∗∗ *P* < 0.001.

#### Net Metabolic Power During Motor Savings

We compared early learning phases across both perturbations and observed main effects of perturbation (*F*(1,28) = 98.6, *P* < 0.001) and interaction of group and perturbation (*F*(1,28) = 4.61, *P* = 0.041) on early savings of net metabolic power, but no group differences (*P* = 0.477) (Fig. 4*C*). Tukey’s comparison showed significant reductions between early adapt 1 and early adapt 2 for both AIH (*t*(28) = 8.54, *P* < 0.001) and control groups (*t*(28) = 5.50, *P* < 0.001). We also observed late savings of net metabolic power between perturbations (*F*(1,28) = 10.8, *P* = 0.003), but no effects of group (*P* = 0.799) nor interaction of group and perturbation (*P* = 0.978). Post-hoc analysis showed magnitude reductions in net metabolic power during late learning in both the AIH (*t*(28) = 2.34, *P* = 0.027) and control groups (*t*(28) = 2.30, *P* = 0.029) (Fig. 4*D*).

#### Correlations of Kinematic, Kinetic, and Metabolic changes

We investigated correlations between changes in net metabolic power and changes in kinematic and kinetic variables. There were non-significant positive relationships between changes in net metabolic power and step width in the fast (R^2^ = 0.203, *P* = 0.092) and slow leg (R^2^ = 0.254, *P* = 0.055). We observed significant correlations between reductions in net metabolic power and decreases in peak ML GRF during the first perturbation, notably during the propulsive phase for the fast leg (R^2^ = 0.728, *P* < 0.001) and during the braking phase for the slow leg (R^2^ = 0.328, *P* = 0.026). These results suggest that individuals with the largest decreases in step width and peak ML GRF drove reductions in net metabolic cost.

## DISCUSSION

This study examined adaptations in step width, peak ML GRF, and their association with changes in net metabolic power during split-belt treadmill walking. We observe distinct coordination strategies between the fast and slow limbs during the initial perturbation exposure. We further demonstrate motor savings of these distinct interlimb adaptations during a subsequent perturbation as well as novel associations between changes in peak ML GRF and reductions in net metabolic power. Repetitive AIH treatments further enhanced both motor learning and motor savings of adaptive frontal plane mechanics.

### Step Width Reductions During Motor Learning and Motor Savings

During adapt 1, we observed that step width increased bilaterally during early learning, followed by a reduction toward baseline as learning progressed. Congruent with our hypothesis, participants utilized an initial compensatory widening of gait to increase stability under the newly altered belt speeds, which was gradually reduced as subjects adapted to the perturbation. Other studies similarly observed initial increases in step width (Fettrow et al., 2021) in addition to increases in ML margin of stability during early learning (Buurke et al., 2018; Brinkerhoff et al., 2024) followed by reductions in step width throughout adapt 1. One novel finding was that the leg on the slow belt followed a similar pattern in foot placement adaptation during a subsequent perturbation, while the fast leg maintained wider steps into late learning. The maintenance of wider steps can likely be interpreted as the fast leg’s strategy to preserve stability (McAndrew Young et al., 2012). These observations indicate that bilateral foot placement adaptations are primarily adopted in the early learning phase of asymmetric walking, whereas re-exposure reveals a distinct shift in fast and slow foot placement strategies. These results contrast with previously reported sagittal plane mechanics during split-belt walking, which shows a return to step length symmetry as adaptation progresses (Bogard et al., 2023; Leech et al., 2018). This suggests that ML spatial adaptations require distinct mechanical strategies between limbs as interlimb symmetry in the frontal plane is not a control priority.

The AIH group uniquely demonstrated early savings of step width in both limbs, indicating that receiving low-oxygen treatments enhances retention of previously learned strategies. These findings support our previous observations that repetitive AIH exposure facilitates improved motor savings of spatiotemporal asymmetry (Bogard et al., 2023). In contrast, we did not observe late savings, indicating that both groups consolidated their adaptation strategies at an earlier timescale regardless of intervention. This interpretation is consistent with sagittal plane spatial adaptations observed in healthy participants (Bogard et al., 2023) and individuals with Parkinson’s disease (Thompson & Reisman, 2022).

### Distinct Interlimb Adaptation Patterns in Peak ML GRF During Motor Learning and Savings

The fast and slow legs differentially adapted peak ML GRF during the braking phase of both perturbation trials. The braking phase of gait, occurring immediately after heel-strike in early stance, is critical for re-stabilization and may explain why higher forces were maintained on the fast leg (Rawal & Singer, 2021). Similarly, our group previously noted larger braking forces on the fast leg in the sagittal plane during split-belt walking (Bogard et al., 2023). In contrast, the leg on the slow belt adapted by decreasing its peak force magnitude, emphasizing each leg’s unique role in maintaining ML stability during split-belt adaptation. Contrary to our findings, Roper et al., 2017 observed no interlimb differences in ML GRF impulses during the braking phase. This discrepancy may arise from the limitations of time integration of force, which might not fully capture key details such as peak magnitudes (Deffeyes & Peters, 2021), particularly in dynamically adaptive environments. One limitation is that we analyzed peak force magnitudes when ML GRF was directed medially, not capturing adaptations of laterally directed forces (John et al., 2012) which could provide further insights into kinetic adaptation strategies.

During the propulsive phase, peak ML GRF of the fast leg reactively increased during early learning before returning to baseline in both perturbation trials. Conversely, peak ML GRF of the slow leg did not alter during adapt 1 but was reduced below baseline during adapt 2. The unique adaptive responses to each belt speed align with previously observed changes in ML GRF at varying walking speeds, as well as magnitude shifts throughout the gait cycle that reflect our observed peak force values during the braking and propulsive phases (John et al., 2012). Although interlimb adaptation patterns during the propulsive phase are distinct, they appear less pronounced compared to the braking phase. Interestingly, the slow leg demonstrated savings of peak ML GRF during the propulsive phase, suggesting that changes during push-off primarily contributed to the observed strong correlation with net metabolic power. Applying external lateral stabilization reduced energetic costs by minimizing excessive ML movement (Dean et al., 2007), while external horizontal aiding forces similarly lowered metabolic rate by offsetting the high energy cost of generating propulsive forces during walking (Gottschall & Kram, 2003). Interestingly, increasing propulsive demand through inclined split-belt walking appears to improve the magnitude of motor learning in the sagittal plane in participants with stroke as evidenced by reductions in step length asymmetry (Sombric & Torres-Oviedo, 2020). This suggests that the kinetic demands of propulsion influence not only impact the energetic cost but also shape the magnitude of motor adaptation. Our observations emphasize the contribution of the ML forces generated during the propulsive phase in metabolic adaptation, underscoring the importance of optimizing push-off mechanics to reduce energy expenditure. Another consideration is that Roper et al. observed that the ML force impulse of the slow limb during late adaptation to be more medially directed than the fast limb. This highlights an important limitation that our analyses of the peak force magnitudes do not adequately capture adaptations in ML GRF directionality and their corresponding effect on net metabolic power (Roper et al., 2017).

We further demonstrated that during re-exposure to the split-belt walking perturbation, individuals in the AIH group uniquely retained motor strategies that reduced peak ML GRF in addition to narrowing step width. Even though both groups similarly demonstrated early savings during the propulsive phase on the fast leg, only the AIH group displayed reduced peak ML GRF on the slow leg during adapt 2. Although we previously observed that anterior-posterior force asymmetry was ultimately reduced during split-belt walking (Bogard et al., 2023), differences between limbs were maintained in frontal plane kinetics during braking and propulsive phases. These distinct adaptations of the fast and slow leg may be partly attributed to independent neural control of each limb, which is modulated based on unique sensory feedback from afferent inputs to each limb (Choi & Bastian, 2007). Furthermore, it has been shown that intralimb adaptations occur at a quicker timescale than interlimb gait parameters (Sato & Choi, 2022), as evidenced by each leg independently adapting its peak ML GRF during adapt 1. Thus, differences in plane-specific adaptations may indicate distinct control priorities where the sagittal plane adaptations prioritize a more symmetrical AP force generation to propel the body forward, while frontal plane adaptations allow asymmetrical stepping mechanics to maintain dynamic stability.

### Energetic Optimization

We observed that both groups concurrently reduced net metabolic power as they adapted their frontal plane mechanics, supporting the notion that adaptive control of dynamic balance is related to the reduction in energy expenditure (Donelan et al., 2001; Finley et al., 2013; Selinger et al., 2015; Buurke et al., 2018). Although the AIH group increased net metabolic power from early to late learning during Adapt 2, they exhibited about 0.3 W/kg less than the controls during early learning and continued to decrease throughout late learning. These observations suggest that the retention of motor adaptations is more pronounced after repetitive AIH, as indicated by the trend in reduced net metabolic power. The savings of frontal plane adaptations align with numerous studies that examined how biomechanical adaptations reduce energetic expenditure upon re-exposure to the perturbation, including decreases in step length asymmetry (Finley et al., 2013; Roemmich & Bastian, 2015; Buurke et al., 2022), optimization of step frequency (Selinger et al., 2015), adjustments in peak force asymmetry (Bogard et al., 2023), and reductions in positive mechanical work (Sánchez et al., 2019). Given previous work that shows savings of biomechanical adaptations are improved with exposure to larger perturbations that utilize great split-belt speed ratios (Leech et al., 2018), we speculate that energetic savings may reach a plateau as split-belt speed ratios increase. The findings of this study, which used a 2:1 speed ratio, contrast with our previous observations using a 1.5:1 speed ratio in which we noted continual reductions in net metabolic power during the subsequent perturbation (Bogard et al., 2023). Nevertheless, both groups demonstrated early savings of lower net metabolic power, with AIH participants demonstrating greater adaptations. Late savings showed similar changes between groups, indicating that gains in metabolic power had stabilized.

### Correlations Between Kinematics, Kinetics, and Net Metabolic Power

Narrowed step widths were positively correlated with reductions in net metabolic power. Other studies have reported reductions in metabolic cost when frontal plane strategies are optimized to walk at preferred step width (Donelan et al., 2001) and exploit frontal plane passive dynamics (Fettrow et al., 2021). In the sagittal plane, reductions in net metabolic power paralleled increases in step time asymmetry (Bogard & Tan, 2024) and decreases in step length asymmetry during motor learning and savings (Sánchez et al., 2019; Bogard et al., 2023). These findings show that the metabolic determinants of split-belt adaptation are driven by distinct yet complementary kinematic adaptations across both planes (Buurke et al., 2020; Buurke & den Otter, 2021) as well as limb-specific kinetics that are likely interdependent. For example, the correlations between ML force production and net metabolic power showed stronger relationships than with kinematic changes. Notably, kinematic adjustments respond more quickly to perturbations and are driven by sensory feedback whereas kinetic adaptations are more gradual (Mawase et al., 2013), supporting the view that ML kinetic demands largely regulate changes in energy cost. There was a significant correlation between ML GRF of the fast leg during the propulsive phase and net metabolic power, suggesting that as the limb transitions from push-off to the swing phase, control of ML GRF may be critical in regulating metabolic cost during gait adaptations. Indeed, modulating propulsive force in the sagittal plane influences walking energetics (Grabowski et al., 2005; Pieper et al., 2021) and our findings suggest that ML mechanics contribute to the overall metabolic cost. Regulation of frontal plane stability is critical in driving initial adaptation, as recent studies show that ML adjustments adapt at a faster timescale than in the sagittal plane (Brinkerhoff et al., 2024). Furthermore, it has been proposed that improvements in sagittal plane symmetry may come at the cost of frontal plane parameters, (Cornwell et al., 2024), a pattern also observed in individuals post-stroke (Buurke et al., 2020). Together with our findings, these observations emphasize the importance of examining adaptations across both planes to gain a comprehensive understanding of how their interactions influence energetic optimization.

### Improved Adaptations Following AIH

We demonstrated enhanced kinematic and kinetic adaptations following repetitive AIH treatments. Notably, the AIH group exhibited early savings of reduced step width for both legs and peak ML GRF during the propulsive phase for the slow leg. The neural mechanisms underlying AIH-induced improvements in motor performance and motor learning remain unclear. Previous studies in spinally injured rats show that BDNF-dependent mechanisms were enhanced following daily AIH, leading to improved horizontal ladder walking (Lovett-Barr et al., 2012). In humans, emerging evidence from our lab and others indicate that AIH strengthens descending neural excitability (Christiansen et al., 2018; Bogard et al., 2023; Bogard, Hembree, et al., 2024) as well as decreases performance fatiguability (Bogard, Pollet, et al., 2024). Together, these findings suggest that similar BDNF-dependent mechanisms underly improvements in motor learning. Indeed, preliminary evidence indicates that BDNF affects other forms of motor learning including visuomotor adaptation and use-dependent plasticity (Fritsch et al., 2010; Joundi et al., 2012; Mang et al., 2014; Helm et al., 2016). We speculate that AIH-induced increases in BDNF enhance synaptic plasticity (Fritsch et al., 2010), leading to improved sensorimotor adaptation (Bogard et al., 2023) and motor savings observed in the present study. Previous research suggests that the nervous system facilitates postural adjustments by sending commands to anticipate and react to postural threats (Cesari et al., 2022). The savings of reductions in step width and ML GRF during adapt 2 support the interpretation of a shift from generalized anticipatory control strategies to task-specific reactive control to preserve stability following a perturbation (Ahuja & Franz, 2022).

Although we did not measure step-to-step adjustments, quantifying any persistent increases step variability throughout adapt 2 may further elucidate shifts towards task-specific reactive control that accommodates flexible foot placement strategies (Ahuja & Franz, 2022). While the current data set cannot parse the underlying neural mechanisms, it remains plausible that AIH-induced synaptic plasticity may enhance these adaptive control mechanisms, leading to more effective dynamic balance during destabilizing perturbations.

AIH treatments have significant potential for integration into the design of rehabilitation training paradigms aimed at improving walking stability and reducing fall risks. Indeed, both long-term and progressive training paradigms have been previously demonstrated to induce improvements in motor learning (Christiansen et al., 2020), lateral balance control (Sawers et al., 2013), and enhanced walking performance after repetitive AIH exposure (Hayes et al., 2014; Tan et al., 2021). Moreover, prolonged exposure to optical flow perturbations has the potential to be used as a training method to improve corrective motor adjustments while walking in older adults (Richards et al., 2019). Improving frontal plane stability is especially critical for older populations and individuals with neurological disorders, considering their diminished motor control and adaptive capabilities (Sato & Choi, 2022; Fettrow et al., 2021; Arora et al., 2020). Therefore, future studies may more critically examine how implementing targeted practice paradigms combined with AIH treatments improves dynamic balance in persons with neurological deficits such as SCI (Navarrete-Opazo et al., 2017) as well as healthy older adults. Our findings further underscore the coupling between adaptive ML control and walking energetics, which may further inform the detection of balance decline as well as the tailoring of patient-specific rehabilitation strategies to improve postural control.

## DATA AVAILABILITY

The authors confirm that the data supporting the findings of this study are fully available and presented in the supporting information of the manuscript. Correspondence and requests for materials should be addressed to A.Q.T.

## GRANTS

This work was funded by the National Institute of Health’s National Center of Neuromodulation for Rehabilitation (NM4R) [NIH P2CHD086844], ABNEXUS Award (AQT), and the Boettcher Foundation Webb Waring Biomedical Research Award (AQT). NMN was supported by The Shurl and Kay Curci Foundation.

## ACKNOWLEDGEMENTS

We thank our study participants and collaborators.

## AUTHOR CONTRIBUTIONS

AQT and ATB designed the study protocol; ATB, AKP, and LP conducted the experiments; ATB, NMN, and LMP analyzed the data; NMN drafted the original manuscript. All authors contributed to the interpretation of the results and revision of the manuscript; All authors approved the final version of the manuscript.

---

**Table 1.**
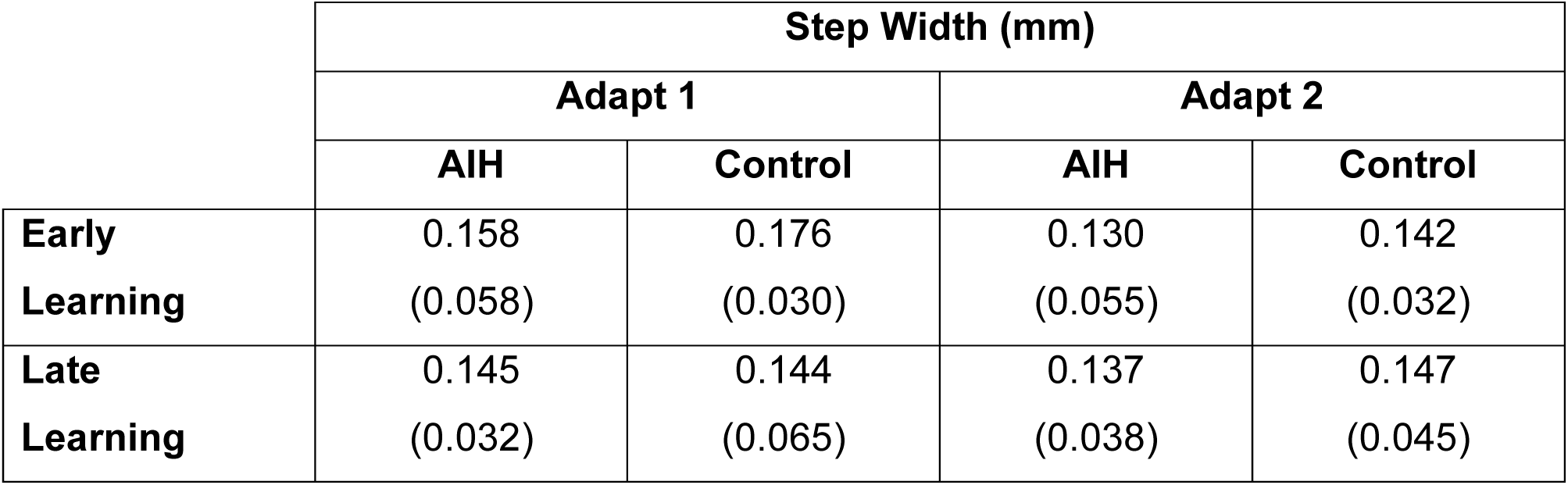
Mean (SD) values of step width (mm) during early and late learning during adapt 1 and adapt 2 for the AIH and Control groups.

**Table 2.** Mean (SD) values of peak ML GRF (BW) during the propulsive and braking phases of early and late learning during adapt 1 and adapt 2 for the AIH and Control groups.

|  |  | Peak ML GRF (BW) |  |  |  |
| --- | --- | --- | --- | --- | --- |
|  |  | Adapt 1 |  | Adapt 2 |  |
|  |  | AIH | Control | AIH | Control |
| **Braking Phase** | **Early Learning** | 0.141<br>(0.046) | 0.123<br>(0.023) | 0.117<br>(0.030) | 0.111<br>(0.024) |
|  | **Late Learning** | 0.114<br>(0.032) | 0.115<br>(0.029) | 0.114<br>(0.028) | 0.111<br>(0.029) |
| **Propulsive Phase** | **Early Learning** | 0.106<br>(0.042) | 0.096<br>(0.034) | 0.078<br>(0.018) | 0.078<br>(0.020) |
|  | **Late Learning** | 0.069<br>(0.015) | 0.073<br>(0.019) | 0.071<br>(0.011) | 0.072<br>(0.014) |

**Table 3.**
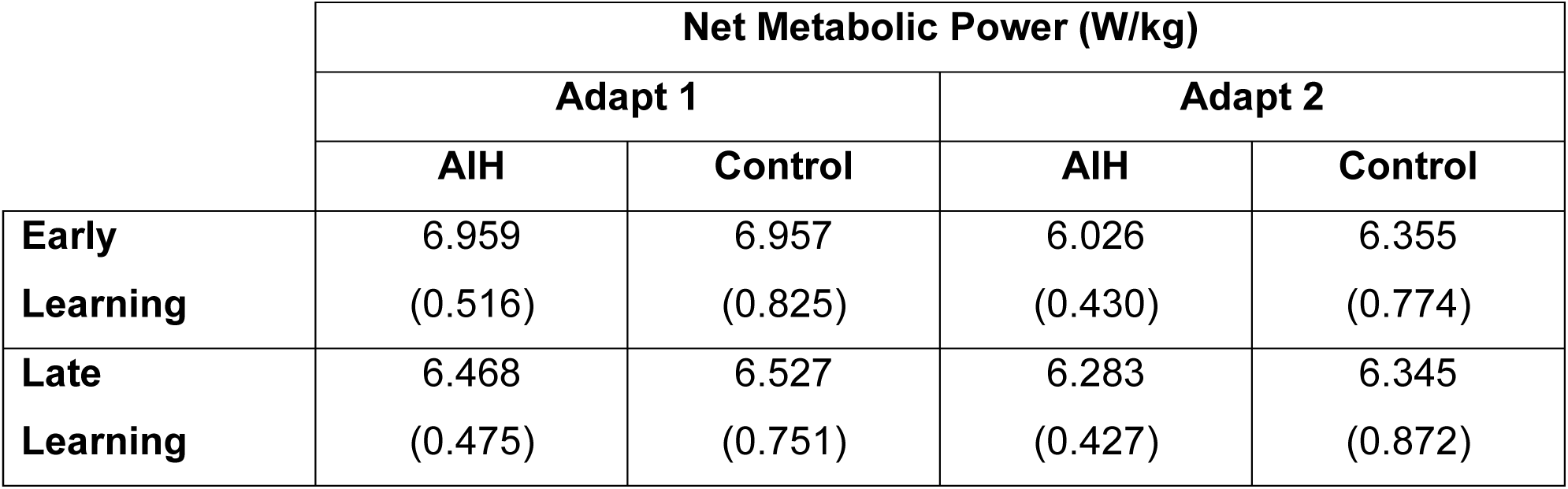
Mean (SD) values of net metabolic power (W/kg) during early and late learning of adapt 1 and adapt 2 for the AIH and Control groups.

|  | Net Metabolic Power (W/kg) |  |  |  |
| --- | --- | --- | --- | --- |
|  | Adapt 1 |  | Adapt 2 |  |
|  | AIH | Control | AIH | Control |
| **Early Learning** | 6.959<br>(0.516) | 6.957<br>(0.825) | 6.026<br>(0.430) | 6.355<br>(0.774) |
| **Late Learning** | 6.468<br>(0.475) | 6.527<br>(0.751) | 6.283<br>(0.427) | 6.345<br>(0.872) |

